# ELAV proteins bind and stabilize C/EBP mRNA in the induction of long-term memory in *Aplysia*

**DOI:** 10.1101/2020.08.14.251389

**Authors:** Anastasios A. Mirisis, Ashley M. Kopec, Thomas J. Carew

## Abstract

Long-term memory (LTM) formation is a critical survival process by which an animal retains information about prior experiences in order to guide future behavior. In the experimentally advantageous marine mollusk *Aplysia*, LTM for sensitization can be induced by the presentation of two aversive shocks to the animal’s tail. Each of these training trials recruits distinct growth factor signaling systems that promote LTM formation. Specifically, whereas intact TrkB signaling during Trial 1 promotes an initial and transient increase of the immediate early gene *apc/ebp* mRNA, a prolonged increase in *apc/ebp* gene expression required for LTM formation requires the addition of TGFβ signaling during Trial 2. Here we explored the molecular mechanisms by which Trial 2 achieves the essential prolonged gene expression of *apc/ebp*. We find that this prolonged gene expression is not dependent on *de novo* transcription, but that *apc/ebp* mRNA synthesized by Trial 1 is post-transcriptionally stabilized by interacting with the RNA-binding protein ApELAV. This interaction is promoted by p38 MAPK activation initiated by TGFβ. We further demonstrate that blocking the interaction of ApELAV with its target mRNA during Trial 2 blocks both the prolonged increase in *apc/ebp* gene expression and the behavioral induction of LTM. Collectively, our findings elucidate both *when* and *how* ELAV proteins are recruited for the stabilization of mRNA in LTM formation.

**Significance Statement:** In the present paper we significantly extend the general field of molecular processing in LTM by describing a novel form of pre-translational processing required for LTM which relies on the stabilization of a newly synthesized mRNA by a unique class of RNA binding proteins (ELAVs). In the broad field of molecular mechanisms of transcription-dependent LTM, there are now compelling data showing that important processing can occur after transcription of a gene, but before translation of the message into protein. Although the potential importance of ELAV proteins in LTM formation has previously been reported, to date there has been no mechanistic insight into the specific actions of ELAV proteins in stabilization of mRNAs known to be critical for LTM. Our new findings thus complement and extend this literature by demonstrating *when* and *how* this post-transcriptional gene regulation is mediated in the induction of LTM.

## Introduction

Long-term memory (LTM) can be mechanistically distinguished from short-term and intermediate-term memory by its dependence on *de novo* transcription (Sutton et al., 2001; Alberini, 2009; Kandel, 2012). LTM for sensitization of the defensive tail and siphon withdrawal responses in *Aplysia californica* provides an experimentally advantageous system in which to study the molecular mechanisms underlying LTM formation. In this system, training trials (brief electrical shocks to the tail) lead to the release of the neuromodulator serotonin (5HT) in the CNS of *Aplysia* (Marinesco and Carew, 2002), which in turn enhances synaptic transmission in the reflex circuit in the pleural-pedal ganglia (Brunelli et al., 1976). Studies focusing on the sensory neurons (SNs) mediating these reflexes have elucidated critical molecular events underlying LTM for sensitization. For example, initiation of CREB-mediated transcription of several target genes, including the immediate early gene *Aplysia c/ebp* (*apc/ebp*), is required for LTM formation (Alberini et al., 1994; Alberini, 2009; Kandel, 2012; Mirisis et al., 2016).

As in many other species, the induction of LTM in *Aplysia* critically depends upon the number and pattern of training trials. For example, two training trials separated by 45 min (but not by 15 or 60 min), induces LTM formation (Philips et al., 2013), revealing a narrow time window during which a second trial interacts with a “molecular context” produced by the first trial to give rise to LTM formation (Philips et al., 2013).

In this paper, we explore the role of mRNA stabilization induced by growth factors (GFs) in LTM formation, with an emphasis on regulation that occurs between transcription and translation within the molecular context established by Trial 1. Kopec et al. (2015) found that signaling through two distinct GF families, TrkB and TGFβ, is required during different training trials for the induction of the molecular events critical for LTM formation. Specifically, *apc/ebp* gene expression induced by Trial 1 peaked at 45 mins, was dependent on TrkB signaling, and returned to baseline by 1 hr (Philips et al., 2013; Kopec et al., 2015). However, if Trial 2 was delivered during peak *apc/ebp* mRNA levels at 45 min after Trial 1, *apc/ebp* mRNA level remained elevated for more than 1 hr, and this elevation critically required TGFβ signaling during Trial 2. Further, the induction of LTM required TrkB signaling during Trial 1 and TGFβ signaling during Trial 2 (Kopec et al., 2015).

Trial 2-dependent *apc/ebp* gene expression was directly dependent on the TrkB-mediated events set in motion by Trial 1 (Kopec et al., 2015). This dependence could be explained in one of two ways: (i) TGFβ signaling in Trial 2 also induces *apc/ebp* gene expression in Trial 2, or (ii) TGFβ signaling in Trial 2 engages post-transcriptional mechanisms which act to stabilize the *apc/ebp* mRNA that was generated by TrkB signaling in Trial 1.

In considering the second alternative, a candidate mechanism for the stabilization of the mRNA is the RNA-binding protein ELAV (embryonic lethal abnormal vision), which confers stability to mRNAs containing AU-rich elements (AREs) in their 3’ UTR (Wang et al., 2000; Brennan and Steitz, 2001). Two ELAV family members, ApELAV1 and ApELAV2, have been identified and cloned in *Aplysia*, and ApELAV1 has been shown to bind to *apc/ebp* mRNA both *in vitro* and *in vivo* (Yim et al., 2006). Whereas ELAV proteins have been implicated in memory formation (Quattrone et al., 2001; Yim et al., 2006; Bolognani et al., 2007), these studies lacked temporal resolution and did not elucidate the mechanisms mediating ELAV-mediated mRNA stabilization in LTM formation.

In the present paper, we examine mRNA stabilization induced by ApELAV during LTM formation and show that it is specifically regulated by TGFβ-induced p38 MAPK signaling initiated by Trial 2. Disrupting the ApELAV-*apc/ebp* interaction blocks the prolonged expression of *apc/ebp* mRNA, and this interaction is required for the behavioral induction of LTM. Collectively, these data demonstrate both *when* and *how* mRNA stabilization is achieved in LTM formation.

## Materials and Methods

### CONTACT FOR REAGENT AND RESOURCE SHARING

Further information and requests for resources and reagents should be directed to and will be fulfilled by the Lead Contact, Thomas J. Carew.

### EXPERIMENTAL MODEL AND SUBJECT DETAILS

*Aplysia californica* were acquired from South Coast Bio-Marine and University of Miami National Resource for *Aplysia.* All animals were allowed to acclimate for at least 4 days in circulating tanks with artificial sea water (ASW) (Instant Ocean) at 15° C.

### METHOD DETAILS

#### Ganglion Preparation

Pleural-pedal ganglia were dissected from anesthetized *Aplysia californica* (150-250 g). In a 1:1 solution of MgCl_2_ and ASW, the pleural SN cluster and SN-MN neuropil were exposed. Ganglia were perfused with ASW for at least 1.5 hr to clear MgCl_2_ prior to experimentation. Experimental and contralateral within-animal control ganglia both received GF/drug/vehicle treatment, while only the experimental ganglia received two-trial 5HT training.

#### Training and Drug Incubation for Gene Expression Analyses

After MgCl_2_ washout, two-trial analog training was delivered by two pulses of 5HT (50 μM, Sigma, #H9523-25MG) in ASW (ITI = 45 min). As previously described (Kopec et al., 2015), ganglia in GF treatment experiments were blocked with BSA for at least 5 min pre-GF treatment, and GF/drug/vehicle was applied at the appropriate experimental time point. Drugs used were as follows: rhTGFβ-1 (R&D Systems, #240-B-002), rhTGFβ-sRII (R&D Systems, #241-R2-025), SB203580 (Tocris, #1202), Actinomycin D (Sigma, #A1410-10MG), CMLD-2 (Millipore, #538339). Drug incubation was from 15 min before the training trial, during the training trial, and proceeded until SN cluster collection. Within-animal control ganglia concurrently received appropriate vehicle but did not receive two-trial training. Control and experimental SN clusters were collected at the experimental time point and prepared for either Western Blotting or quantitative PCR analysis.

#### ApELAV1 Protein Expression

The coding region for ApELAV1 was isolated by PCR from a previously published plasmid kindly provided by the Bong-Kiun Kaang (Seoul, South Korea)(Yim et al., 2006) and ligated into pGEX vector to make pGEX-ApELAV1-GST. Rosetta 2 DE3 cells (Millipore) were transformed with this plasmid, and cells were induced for 2 h at 37°C with 1 mM IPTG during log-phase growth. Cells were subsequently lysed and subjected to Western blotting.

#### Western Blotting

Samples were lysed and snap-frozen in RIPA buffer (Sigma) with 1X Halt protease and phosphatase inhibitor cocktail (Invitrogen), loaded onto 4-12% Bis-Tris gels (Novex; Life Technologies) for electrophoresis and then transferred to nitrocellulose membranes for incubation with primary antibodies and secondary antibodies: anti-HuR (Santa Cruz, sc-5261), anti-β-Actin (Cell Signaling Technology, #3700), anti-p-p38 MAPK (Cell Signaling Technology, #4511), anti-β-Tubulin (Cell Signaling Technology, #2146), anti-mouse IgG IRDye 680LT (LICOR Biosciences, #926-68020), anti-rabbit IgG IRDye 800CW (LICOR Biosciences, #926-32211). For ApELAV and p-p38 MAPK experiments, protein levels within each sample was assessed using the LICOR imaging system. Each protein of interest band was normalized to the tubulin or actin within the same sample, and normalized protein of interest in the experimental ganglia was compared to within-animal control ganglia. Data are displayed as fold-expression of normalized protein of interest in experimental sample relative to control sample.

#### RNA isolation, cDNA synthesis, quantitative PCR analysis

RNA was isolated and purified using Ambion RNAqueous Micro kits (Invitrogen), and total RNA for cDNA synthesis was normalized between experimental and control ganglia (~100ng RNA). cDNA was synthesized using Superscript IV cDNA synthesis reagents (Invitrogen). Quantitative PCR was performed using a Roche LightCycler 480, with 1 μL cDNA, 1 μL each of 5μM F’ and R’ primers, and 18 μL 1X SybrGreen (Roche). Primer sequences are as follows: apc/ebp-F: caccacctcactcccatctc; apc/ebp-R: ctgacgtctgcgagactttg; apgapdh-F: ctctgagggtgctttgaagg; apgapdh-R: gttgtcgttgagggcaattc. The amplification program was 95°C for 3 mins, 30 cycles of 95°C for 10 s, 60°C for 20 s, and 72°C for 30 s, and lastly a melting curve to confirm a single PCR product. Data were analyzed by normalizing the quantity of each gene of interest (measured by Ct) to *apgapdh* within the same sample (ΔCt), then comparing normalized values between experimental and within-animal control groups (ΔΔCt). Data are displayed as fold induction relative to the control group.

#### Single molecule fluorescence *in situ* hybridization (smFISH)

*Aplysia* SNs were isolated from the ventral SN cluster of the pleural ganglion of 80g animals according to (Zhao et al., 2009) in 50mm Mattek plates. After 4 days *in vitro*, 7 plates of 10-20 SNs were chosen and separated into 4 different groups randomly. 2 plates each were designated as “2X 5HT +DMSO”, “2X 5HT +SB”, and “2X ASW +DMSO”. 1 plate received no treatments but was used as a “no probe” control to determine background fluorescence of SNs without DNA probes.

The smFISH protocol was adapted from (Eliscovich et al., 2017). Following treatment according to the experimental paradigm (5HT treatment was 50 μM in ASW, ASW treatment was an equivalent volume of ASW), the cells were fixed for 30 min at room temperature in 4% formaldehyde in 30% sucrose / DEPC-treated PBS with supplemented 1 mM magnesium chloride and 0.1 mM calcium chloride (DEPC-PBS-MC). Cells were washed quickly 2X with DEPC-PBS-MC, and quenched with glycine in PBS-MC. Quench buffer was removed and cells were incubated in permeabilization-block buffer (0.2% Triton X-100, 0.5% Ultrapure BSA in DEPC-PBS-MC) for 30 min at room temperature while shaking. Permabilization-block buffer was removed and cells were incubated with pre-hybridization buffer (10% deionized formamide, 2X SSC, 0.5% Ultrapure BSA in nuclease-free water) for 30 min at room temperature without shaking. Pre-hybridization buffer was removed and cells were incubated with hybridization buffer (10% deionized formamide, 2X SSC, 1 mg/mL *Escherichia coli* tRNA, 20 mg/mL BSA, 2X SSC, 2 mM Vanadyl Ribonucleoside Complex, 10% dextrane sulfate, 125 nM of DNA probes in nuclease-free water) for 4 h at 37 °C. Thirty-three (33) DNA probes conjugated to Quasar 670 dye (LGC Biosearch Technology) were designed to be antisense to all regions on *apc/ebp* mRNA without significant complementarity to other mRNAs in the *Aplysia* Refseq database (NCBI) and are listed in Supplementary Table 1. Cells were washed four times with 2X SSC and then once quickly with nuclease-free water. Coverslips were detached from the plates and mounted using Prolong Gold Antifade Reagent with DAPI (Life Technologies).

A Deltavision Elite imaging system (GE) equipped with Olympus ×60 / 1.42 NA oil immersion lens and an InsightSSI Solid State Illumination module was used to image samples. Individual full SN somata were imaged in a z-stack with 500 nm spacing. Raw images were loaded into FISH-QUANT software (Mueller et al., 2013) and batch processed after detecting nucleus size (by DAPI) and tracing the outline of each SN. Initial analyses determined that, in the cells which received sham treatment (2X ASW), the number of puncta counted in each cell was correlated strongly to the area of the cell nucleus (Supplementary Figure 3), which corresponds to previous literature noting that the absolute amount of RNA can be dependent on cell size (Marguerat and Bähler, 2012). Therefore, the number of puncta counted in each cell was normalized to the area of the nucleus in each cell, as this value was calculated automatically by FISH-QUANT.

#### Cross-Linking Immunoprecipitation-qPCR

The protocol was adapted from (Yoon and Gorospe, 2016). Experimental and contralateral within-animal control ganglia both received GF chimera/drug/vehicle treatment, while only the experimental ganglia received two-trial 5HT training. At the appropriate experimental timepoint, experimental and control ganglia were exposed twice to 400 mJ UV (254nm) to crosslink ribonucleoprotein complexes. SN clusters of 8 animals were collected and lysed in ice-cold NP-40 lysis buffer (20 mM Tris-HCl pH 7.5, 100 mM KCl, 5 mM MgCl_2_, 0.5% NP-40, 1 mM DTT) supplemented with 1X Halt protease and phosphatase inhibitor cocktail and 40 U/mL RNaseOUT (Invitrogen), and pooled for each experimental and contralateral control sample. Lysates were split into two fractions each and immunoprecipitated with either antibodies against HuR (Santa-Cruz, sc-5261) or normal mouse IgG1 (Santa Cruz, sc-3877) using Protein G Dynabeads (Invitrogen). Beads were washed with ice-cold NT-2 buffer (50 mM Tris-HCl pH 7.5, 150 mM NaCl, 1 mM MgCl_2_, 0.05% NP-40). Following consecutive digestions with DNase I and Proteinase K (New England Biolabs), RNA was isolated with Trizol extraction (Invitrogen) and cDNA was synthesized using Superscript IV cDNA synthesis reagents (Invitrogen). Quantitative PCR was performed as described above. Importantly, primers were designed to lie within the 3’ UTR of *apc/ebp* mRNA, the binding region of ELAV. The primer sequences were as follows: apc/ebpUTR-F: cccaagcatgttgtgatagttgt; apc/ebpUTR-R: tgggaggagatagaagcagtg. Data were analyzed by normalizing the difference in *apc/ebp* between ELAV- and IgG-immunoprecipitated samples to the difference in *apgapdh* between the same samples, then comparing these normalized values between experimental and within-animal control groups.

#### Semi-intact Behavioral Preparation

Preparations were completed as previously described (Sutton et al., 2001; Kopec et al., 2015). Briefly, the ring ganglia (cerebral and paired pleural-pedal ganglia) from anesthetized Aplysia (250-400 g; South Coast Bio-Marine) were surgically isolated, leaving p9 and pleural-abdominal innervation to the tail and abdominal ganglia, respectively, intact. The tail and mantle were surgically removed, and the siphon artery was cannulated with Dow Corning silastic tubing (0.025 in I.D.; Fisher Scientific) and perfused at ~5 ml/min, while the tail was perfused at ~0.5 ml/min via three 22-gauge needles inserted into the tail. The tail and mantle were pinned to the chamber floor, while the ring ganglia with both pleural-pedal ganglia de-sheathed were pinned in an isolated CNS chamber. Both the chamber containing the peripheral tissues as well as the CNS chamber were continuously perfused with seawater (Instant Ocean, 15°C). The p9 and pleural-abdominal nerves exited the CNS chamber through small slits that were sealed with Vaseline. Preparations were allowed at least 2 hrs to recover prior to baseline measurements.

#### Training and Drug Incubation for LTM Analyses

An average of 3-4 baseline siphon-withdrawal measurements were recorded by stimulating the medial posterior tip of the tail with a water jet (0.4 s, 45 psi; ITI 15 mins; Teledyne Water Pik) and measuring the duration of time with which the siphon withdrew. No significant differences in the pre-training baseline duration of T-SWRs were observed in any experimental condition. The CNS was incubated with Drug/Vehicle 15 mins prior to Trial 2, which was washed out 30 mins after Trial 2. Two-Trial LTM training (100 mA, 1.5 s, ITI 45 mins) was delivered medially to the anterior portion of the tail through a hand-held electrode. LTM for sensitization of the T-SWR was assessed by 3-4 tests (ITI=15 mins) 15-22 hrs after training by an experimenter blind to the experimental condition (Drug vs. Vehicle). The data are displayed as duration of siphon withdrawal as a percent of pre-training baseline.

### QUANTIFICATION AND STATISTICAL ANALYSIS

Data were analyzed with parametric statistics using GraphPad Prism. Statistical details for all experiments can be found within the main text and/or in the figure legends. Within-group analyses were performed using paired t-test. LTM for sensitization of the T-SWR is compared relative to pre-training baseline T-SWR within the same preparation. Between-group analyses (Drug vs. Vehicle) were performed using t-tests. When appropriate, ANOVA analyses were used to determine if there was a difference among the groups and are indicated in the main text. Significant results were followed by planned parametric comparisons. Outliers greater than 2 standard deviations from the mean were excluded resulting in the removal of 5 data points (<5% of all data points; reported in Table 1). Due to normal distribution of data, data in all figures are depicted as means ± SEM. Significant within-group comparisons are displayed as asterisks above the summary data, and significant between-group comparisons are displayed as asterisks above a bar drawn between the summary data. Asterisks indicate statistical significance (\**p*<0.05, \*\**p*<0.01).

**Table 1.** Statistical table. Refer to Methods for information on statistics.

|  | Corresponding Figure | Test | Statistics |
| --- | --- | --- | --- |
| A | 1A | Within-group paired *t*-test | -ActinoD: $n=8$, $1.456 \pm 0.140$, $p<0.05$ |
| B | 1A | Within-group paired *t*-test | +ActinoD: $n=7$, $0.8505 \pm 0.1601$ |
| C | 1A | Between-group unpaired *t*-test | Control versus Actino: $t=2.862$, $df=13$, $p<0.05$ |
| D | 1B | Within-group paired *t*-test | 1X 5HT -ActinoD: $0.9600 \pm 0.1118$, $n=7$, $p=0.73$; 2X 5HT -ActinoD: $1.765 \pm 0.1463$, $n=11$, $p<0.01$; 2X 5HT +ActinoD: $1.527 \pm 0.1830$, $n=8$, $p<0.05$ |
| E | 1B | ANOVA | $F=6.786$; $p<0.01$ |
| F | 1B | Between-group unpaired *t*-test | 1X 5HT -ActinoD versus 2X 5HT -ActinoD: $t=3.926$, $df=16$, $p<0.01$; 1X 5HT -ActinoD versus 2X 5HT +ActinoD: $t=2.550$, $df=13$, $p<0.05$; 2X 5HT -ActinoD versus 2X 5HT +ActinoD: $t=1.028$, $df=17$, $p=0.32$ |
| G | 1C | Within-group paired *t*-test | 1X 5HT +TGFβ: $1.384 \pm 0.1493$, $n=8$, $p<0.05$ |
| H | 1C | Between-group unpaired *t*-test | 1X 5HT -ActinoD versus 1X 5HT +TGFβ: $t=2.219$, $df=13$, $p<0.05$ |
| I | 1C | Within-group paired *t*-test; Between-group unpaired *t*-test | 1X 5HT +TGFβ +Actino: $1.230 \pm 0.0954$, $n=12$, $p<0.05$; 1X 5HT +TGFβ versus 1X 5HT +TGFβ +Actino: $t=0.9160$, $df=18$, $p=0.37$ |
| J | 2A | Within-group paired *t*-test | 2X 5HT: $1.845 \pm 0.232$, $n=6$, $p<0.05$ |
| K | 2A | Within-group paired *t*-test; Between-group unpaired *t*-test | 1X 5HT: $1.089 \pm 0.184$, $n=7$, $p=0.99$; 2X 5HT +TGFβsRII: $1.148 \pm 0.086$, $n=7$, $p=0.25$; 1X 5HT versus 2X 5HT: $t=2.582$, $df=11$, $p<0.05$; 2X 5HT versus 2X 5HT +TGFβsRII: $t=2.996$, $df=11$, $p<0.05$ |
| L | 2B | ANOVA | $F=7.804$, $p<0.01$ |
| M | 2B | Within-group paired *t*-test; Between-group unpaired *t*-test | 2X 5HT +DMSO: $186.9\% \pm 23.85\%$, $n=23$, $p<0.01$, 1 outlier; 2X 5HT +SB: $94.82\% \pm 15.74\%$, $n=15$, $p=0.747$, 2 outliers; 2X ASW +DMSO: $100\% \pm 12.74\%$, $n=23$, 1 outlier; 2X 5HT +DMSO versus 2X 5HT +SB: $t=2.855$, $p<0.01$; 2X 5HT +DMSO versus 2X ASW +DMSO: $t=3.213$, $p<0.01$ |
| N | 2C | Within-group paired *t*-test; Between-group unpaired *t*-test | 2X 5HT +SB: $0.734 \pm 0.161$, $n=8$, $p=0.142$; 2X 5HT versus 2X 5HT +SB: $t=4.692$, $df=17$, $p<0.01$ |
| O | 2C | Within-group paired *t*-test; Between-group unpaired *t*-test | 2X 5HT +CMLD-2: $1.194 \pm 0.201$, $n=9$, $p=0.62$; 2X 5HT versus 2X 5HT +CMLD-2: $t=2.350$, $df=18$, $p<0.05$ |
| P | 2C | Within-group paired *t*-test | 1X 5HT +TGFβ +CMLD-2: $0.9825 \pm 0.1928$, $n=7$, $p=0.93$, 1 outlier: see Methods |
| Q | 3A | Within-group paired *t*-test | 2X 5HT: $201.6\% \pm 19.89$, $n=5$, $p<0.01$ |
| R | 3A | Within-group paired *t*-test; Between-group unpaired *t*-test | 2X 5HT +TGFβsRII: $119.0\% \pm 10.31$, $n=3$, $p=0.21$; 2X 5HT versus 2X 5HT +TGFβsRII: $t=2.995$, $df=6$, $p<0.05$ |
| S | 3A | Within-group paired *t*-test; Between-group unpaired *t*-test | 2X 5HT +SB: $124.0\% \pm 18.35$, $n=3$, $p=0.32$; 2X 5HT versus 2X 5HT +SB: $t=2.611$, $df=6$, $p<0.05$ |
| T | 3A | Within-group paired *t*-test; Between-group unpaired *t*-test | 2X 5HT +CMLD-2: $104.4\% \pm 7.91$, $n=3$, $p=0.63$; 2X 5HT versus 2X 5HT +CMLD-2: $t=3.579$, $p<0.05$ |
| U | 4A | Within-group paired *t*-test; Between-group unpaired *t*-test | Vehicle: $128.5\% \pm 7.294$, $n=8$, $p<0.05$; CMLD-2: $109.3\% \pm 6.008$, $n=8$, $p=0.33$, Vehicle versus CMLD-2: $t=2.032$, $df=14$, $p<0.05$ |

## Results

### The Trial 1-dependent increase in *apc/ebp* mRNA level is transcription-dependent

We have shown that *apc/ebp* gene expression is increased 45 min following Trial 1 (Philips et al., 2013; Kopec et al., 2015). Thus, we first asked whether Trial 1-dependent increase in *apc/ebp* gene expression was due to (i) increased gene transcription, or (ii) alternative mechanisms, such as an increase in transcript stability of baseline levels of *apc/ebp* mRNA.

In order to distinguish between these possibilities, dissected pleural-pedal ganglia were incubated with the irreversible transcription inhibitor Actinomycin D (40 μg/mL, Sigma) for 15 min, and contralateral within-animal control ganglia were incubated with vehicle (an equivalent volume of ddH_2_O). Following drug washout, experimental ganglia were exposed to 5HT (50 μM in ASW, Sigma) for 5 min (Trial 1) to simulate a tail shock training trial (Philips et al., 2013). 5HT was washed out with fresh ASW, and pleural SN clusters were collected at 45 min and analyzed for *apc/ebp* gene expression via quantitative PCR. In this and all following experiments, data from drug conditions are expressed as a ratio to vehicle conditions. Consistent with previous results (Kopec et al., 2015), in experimental ganglia which had not received inhibitor pre-treatment, a within-group comparison revealed a significant increase in *apc/ebp* gene expression in SN clusters collected 45 min after Trial 1 (Figure 1a; Table 1A). No significant difference in experimental (+ActinoD) and within-animal control samples (contralateral SN clusters) was observed (Figure 1a; Table 1B). A between-group comparison revealed a significant difference between *apc/ebp* gene expression of SN clusters which had received drug treatment compared to SN clusters which had not received drug treatment (Figure 1a; Table 1C), demonstrating that Trial 1-dependent *apc/ebp* gene expression at 45 min after Trial 1 is dependent on *de novo* gene transcription.

**Figure 1.**
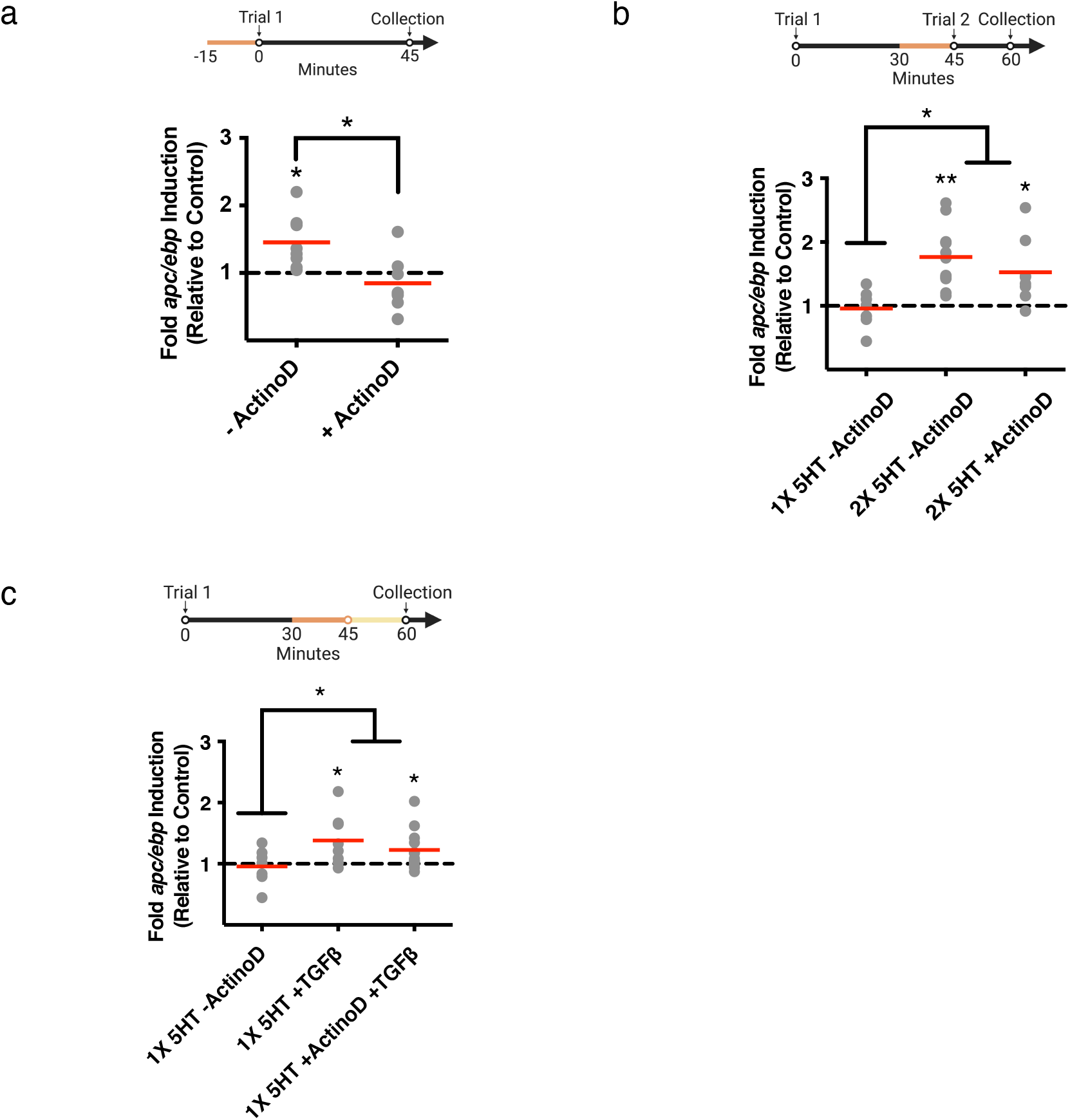
Trial 1 initiates a transcription-dependent increase in *apc/ebp* mRNA levels, whereas Trial 2 prolongs this gene expression through a transcription-independent mechanism. **(a) (Top)** Experimental paradigm. Trial 1 is delivered at time=0:00 and SN somata are collected at 45 min following the onset of Trial 1. Dissected ganglia are treated with actinomycin D 15 min before Trial 1. Drug conditions are expressed as a ratio to vehicle conditions. **(Bottom)** Blocking transcription before Trial 1 significantly disrupts *apc/ebp* gene expression at 45 min. Histograms are labeled by their corresponding drug condition. In this and all subsequent figures, unless specified otherwise, data are displayed as aligned dot plots. Means are represented as red lines within each histogram. Within-group statistical significance is displayed with an asterisk above histograms, and between-group statistical significance is indicated above the histograms being compared. *p<0.05; n=7-8 **(b) (Top)** Experimental paradigm. SN somata are collected at 1 hr following the onset of Trial 1. Dissected ganglia are treated with actinomycin D 15 min before Trial 2. Drug conditions were expressed as a ratio to vehicle conditions. **(Bottom)** A second training trial (Trial 2) at 45 min following the onset of Trial 1 results in a significant transcription-independent significant increase in *apc/ebp* gene expression at 1 hr. \**p*<0.05, \*\**p*<0.01; *n*=7-11. **(c) (Top)** Experimental paradigm. Trial 1 is delivered and SN somata are collected at 1 hr following the onset of Trial 1. Dissected ganglia are treated with actinomycin D or vehicle 15 min before washout (light yellow line) and TGFβ-1 or vehicle is applied at 45 min after Trial 1 until collection. Drug conditions were expressed as a ratio to vehicle conditions. **(Bottom)**Treatment with TGFβ-1 alone at 45 min results in a transcription-independent increase in *apc/ebp* mRNA level. \**p*<0.05; *n*=7-12

### The Trial 2-dependent increase in *apc/ebp* mRNA level is transcription-independent

When Trial 2 is delivered 45 min after Trial 1, *apc/ebp* gene expression at 1 hr after Trial 1 is dependent on (i) TrkB signaling during Trial 1, and (ii) TGFβ signaling during Trial 2 (Kopec et al., 2015). This suggests that either (i) Trial 2 TGFβ-dependent mechanisms cause a new wave of *apc/ebp* transcription, or (ii) Trial 2 induces a form of transcription-independent stabilization of *apc/ebp* mRNA synthesized from Trial 1 (via TrkB-dependent mechanisms). Ganglia were treated with Actinomycin D for 15 min before Trial 2. Following drug washout, experimental ganglia were exposed to 5HT for 5 min (Trial 2). 5HT was washed out with fresh ASW, and pleural SN clusters were collected at 1 hr after Trial 1. A within-group analysis revealed a significant increase in *apc/ebp* mRNA levels in ganglia which had received Actinomycin D (Figure 1b; Table 1D). ANOVA revealed a significant difference among the three groups (Table 1E). Subsequent planned t-tests revealed a significant increase in *apc/ebp* mRNA levels in both the 2X 5HT −ActinoD and 2X 5HT +ActinoD groups, compared to ganglia that had received only 1X 5HT, and no significant difference was noted between the 2X 5HT −ActinoD and 2X 5HT +ActinoD groups (Figure 1b; Table 1F). These data demonstrate that the Trial 2-dependent increase in *apc/ebp* mRNA level at 1 hr after Trial 1 is not dependent on *de novo* gene transcription and supports the hypothesis that the increased *apc/ebp* mRNA level may result from TGFβ-dependent mRNA stabilization.

### TGFβ-1 treatment at 45 min is sufficient for Trial 2-dependent increase in*apc/ebp* mRNA level

By sequestering endogenously released TGFβ-like ligands during Trial 2, we have previously shown that intact TGFβ signaling is required during Trial 2 for *apc/ebp* gene expression at 1 hr (Kopec et al., 2015). We here sought to determine whether treatment with recombinant human TGFβ-1 alone could duplicate these effects.

To investigate this question, ganglia were treated with TGFβ-1 (100 ng/mL, R&D Systems) starting at 45 min after Trial 1 until SN cluster collection and lysis at 1 hr after Trial 1. Treatment with TGFβ-1 has previously been demonstrated to (i) recruit persistent ERK signaling, a necessary step in LTM formation, and (ii) induce long-term facilitation (LTF) in isolated *Aplysia* ganglia (Zhang et al., 1997; Shobe et al., 2016). Contralateral control ganglia were treated with an equivalent volume of vehicle (0.1% BSA in ASW). Treatment with TGFβ-1 resulted in a significant increase in *apc/ebp* mRNA levels when compared to a within-animal control of SN clusters that had received only vehicle (Figure 1c; Table 1G). Planned comparisons further revealed a significant difference between the Trial 1 only and TGFβ-treated groups (Figure 1c; Table 1H).

Given that TGFβ-1 treatment (in the absence of 5HT) at 45 min after Trial 1 is sufficient to increase *apc/ebp* mRNA level at 1 hr (Figure 1c), we further asked whether the TGFβ signaling-dependent effect of increased *apc/ebp* mRNA level at 1 hr after Trial 1 required transcription. Ganglia were treated at 30 min after Trial 1 with Actinomycin D. At 45 min after Trial 1, Actinomycin D was washed out and ganglia were immediately treated with TGFβ-1 until SN cluster collection and lysis at 1 hr after Trial 1. Contralateral within-animal control ganglia received vehicle (0.1% BSA in ASW and ddH2O). TGFβ-1 treatment concomitant with a transcriptional block still resulted in significantly increased *apc/ebp* mRNA level when compared to within-animal control SN clusters. Moreover, there were no significant differences between groups which received TGFβ treatment alone and TGFβ treatment with ActinoD (Figure 1c: Table 1I). Collectively, these data support the hypothesis that TGFβ signaling during Trial 2 is responsible for the post-transcriptional stabilization effect on *apc/ebp* mRNA.

### p38 MAPK is activated by two-trial training and is blocked by blocking TGFβ signaling

Since ELAV is known to be an RNA-binding protein yielding a stabilizing effect on its target mRNAs (Brennan and Steitz, 2001; Yim et al., 2006), we explored the possibility that *apc/ebp* gene expression is modulated by ApELAV. How might ELAV be recruited to exert its stabilizing effects on target mRNAs? First, ELAV protein levels could be increased, or second, ELAV protein could be covalently modified, which would promote its ability to bind its target transcripts.

To distinguish between these possibilities, we first asked whether the protein levels of ApELAV were regulated by two-trial training. To assay changes in protein levels, we used an antibody against HuR, a mammalian member of the ELAV family. This antibody recognizes both *Aplysia* ELAV family members, ApELAV1 and ApELAV2 (Supplementary Figure 1a).

At the 1 hr timepoint (15 min after the onset of Trial 2), pleural SN clusters were collected and subjected to SDS-PAGE and Western blotting. Our results revealed no difference in ApELAV protein levels between the experimental and within-animal control ganglia, which had not received two-trial training (Supplementary Figure 1b_2_).

These findings, however, do not preclude the possibility of a post-translational modification of existing ApELAV proteins (such as phosphorylation) downstream of TGFβ signaling, which would result in an increase of ApELAV-*apc/ebp* mRNA association. In support of this possibility, a particular kinase, p38 MAPK, is both activated downstream of TGFβ signaling (Yamashita et al., 2008), and has previously been identified as a critical mediator of ELAV activation by phosphorylation(Pascale et al., 2005; Lafarga et al., 2009; Bai et al., 2012; Eberhardt et al., 2012; Slone et al., 2016). Our experiments revealed a significant increase in p38 MAPK activation following two-trial training (Figure 2a: Table 1J). This increase was blocked either (i) by eliminating Trial 2, or (ii) by delivering Trial 2 in the presence of TGFβsRII (5 μg/mL, R&D Systems), which acts as an inhibitor of TGFβ signaling by sequestering extracellular ligands (Figure 2a; Table 1K). These data demonstrate that p38 MAPK activation at 1 hr is dependent on TGFβ signaling and may serve an intermediate role between TGFβ signaling and ELAV activation.

**Figure 2.**
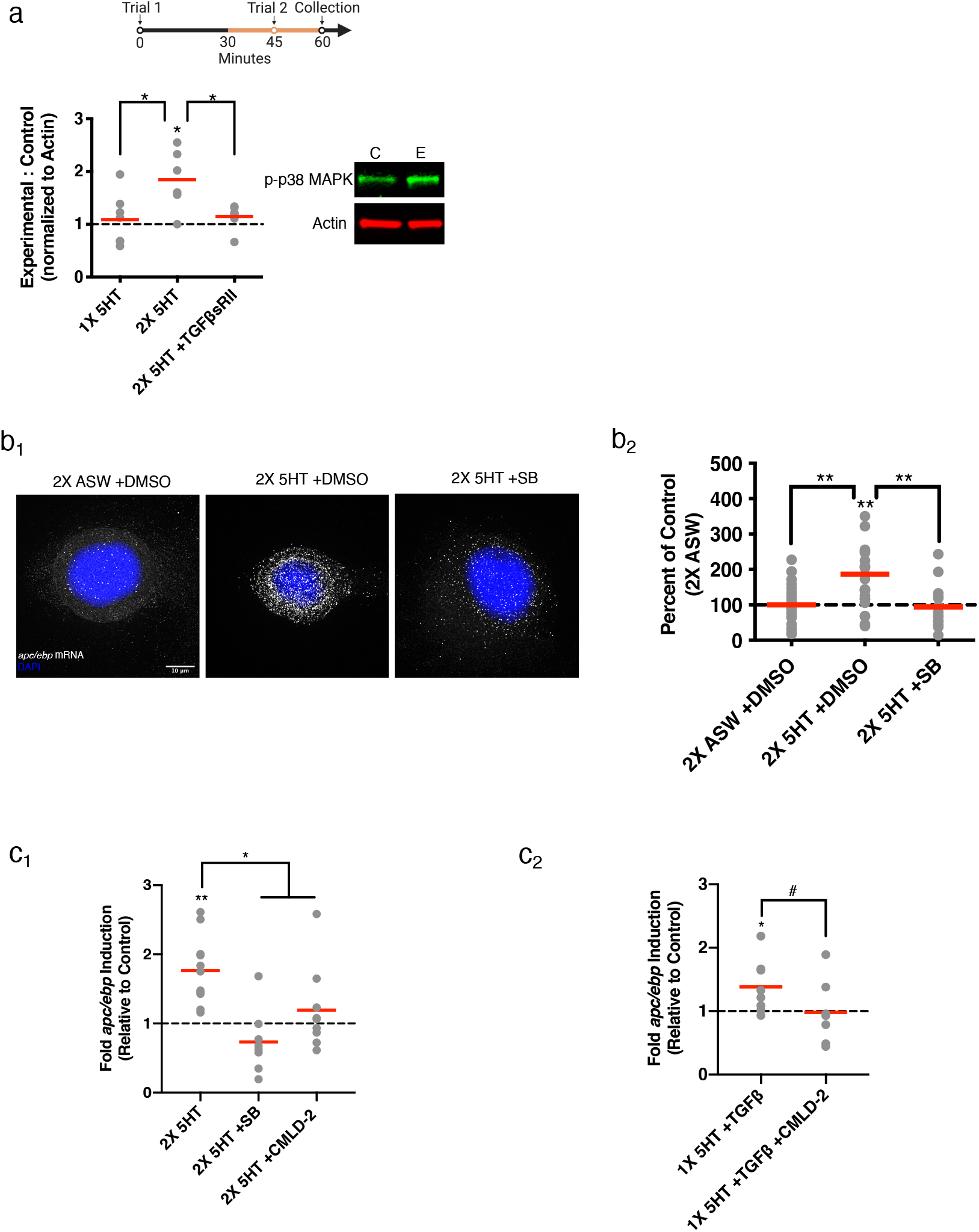
ELAV-mediated prolonged expression of *apc/ebp* mRNA requires p38 MAPK activity initiated by TGFβ signaling. **(a) (Top)** Experimental paradigm. In all paradigms, SN somata are collected at 1 hr following the onset of Trial 1. Dissected ganglia are treated with CMLD-2, SB203580, or TGFβsRII from 15 min before Trial 2 until collection. Dissected ganglia are treated with TGFβ-1 at 45 min until collection. Dissected ganglia are treated with CMLD-2 from 15 min before TGFβ-1 application until collection. Drug conditions were expressed as a ratio to vehicle conditions. **(Bottom)** Following two-trial training, there is a significant increase in p38 MAPK activation which is blocked by treatment with TGFβsRII. \**p*<0.05; n=6-7 **(b_1_)** Representative maximum projection images of single isolated SNs. **(b_2_)** Puncta abundance in isolated SNs. Data are represented as a percentage of puncta abundance in SNs which received two-trial training in the presence of vehicle without 5HT. **p<0.01; n=19-23 **(c_1_)**Blocking the ApELAV-*apc/ebp* mRNA interaction significantly disrupts the increase in *apc/ebp* mRNA level at 1 hr after Trial 1. Inhibiting p38 MAPK activity similarly disrupts the increase in *apc/ebp* mRNA level. \**p*<0.05, \*\**p*<0.01; *n*=7-11. **(c_2_)** Blocking the ApELAV-mRNA interaction blocks the increase in *apc/ebp* mRNA level by TGFβ #*p*<0.3, \**p*<0.05; n=7-8

### p38 MAPK activity is required for prolonged increase in *apc/ebp* mRNA levels

To further examine the role of p38 MAPK in the stabilization of *apc/ebp* mRNA, we directly measured *apc/ebp* mRNA levels, by performing single molecule fluorescence *in situ* hybridization (smFISH) on isolated *Aplysia* SNs (Figure 2b). This technique allows for fluorescent visualization of individual *apc/ebp* RNA molecules as puncta resulting from the combined fluorescence of 33 antisense DNA probes (LGC Biosearch Technology). ANOVA revealed a significant difference in puncta abundance between: (i) SNs that received two-trial training in the presence of vehicle, (ii) SNs which received two-trial training in the presence of SB203580, a p38 MAPK inhibitor previously used and characterized in *Aplysia* (Zhang et al., 2017) (10 μM, Tocris), and (iii) SNs that received two-trial training without 5HT and in the presence of vehicle (Figure 2b_2_; Table 1L). Planned t-tests revealed a significant increase in puncta in SNs that were treated with two-trial training with 5HT in the presence of DMSO compared to each of the other groups (Figure 2b_2_; Table 1M).

In order to examine whether p38 MAPK activation is important for the increase in *apc/ebp* mRNA level 1 hr after Trial 1, ganglia received Trial 2 training with 5HT in the presence of SB203580. 5HT was washed out following Trial 2, and SB203580 treatment proceeded until SN cluster collection and lysis at 1 hr. Within-animal control ganglia were treated with vehicle (0.1% DMSO) but did not receive two-trial training. Treatment with SB203580 indeed resulted in a significant block of the Trial 2-dependent increase in *apc/ebp* mRNA level when compared to two-trial training alone (Figure 2c_1_; Table 1N).

### ApELAV-mRNA interaction is required for prolonged increase in *apc/ebp* mRNA levels

In order to directly manipulate the ability of ApELAV to interact with its target transcripts, we used CMLD-2 (100 μM, Millipore), a potent inhibitor of HuR-ARE interaction (Wu et al., 2015; Slone et al., 2016; Muralidharan et al., 2017). HuR is a mammalian homolog of ELAV, and also contains three highly conserved RNA recognition motifs which facilitate its binding to AREs of mRNAs. Since CMLD-2 directly binds to the highly evolutionary conserved RNA-binding pocket of HuR, we hypothesized that it would likewise bind and inhibit ApELAV binding. We found that following treatment of ganglia with Actinomycin D to inhibit basal *de novo* transcription of *apc/ebp*, addition of CMLD-2 accelerates *apc/ebp* mRNA degradation compared to control (0.8% acetonitrile and ddH_2_O) (Supplementary Figure 2a_2_).

In order to examine whether the ApELAV-*apc/ebp* mRNA interaction plays a critical role in increased *apc/ebp* mRNA levels during the two-trial training paradigm, ganglia received Trial 2 in the presence of CMLD-2. 5HT was washed out following Trial 2, and CMLD-2 treatment proceeded until SN cluster collection and lysis at 1 hr. Within-animal control ganglia were treated with vehicle but did not receive two-trial training. We found that treatment with CMLD-2 significant blocked the increase of Trial 2-dependent *apc/ebp* mRNA levels when compared to two-trial training alone (Figure 2c_1_; Table 1O). Treatment with vehicle alone did not affect the Trial 2-dependent increase in *apc/ebp* mRNA level (data not shown). A similar effect on *apc/ebp* mRNA level was observed when CMLD-2 was co-administered with TGFβ-1 treatment in lieu of 5HT. No significant difference between experimental and within-animal control ganglia were observed (Figure 2c_2_; Table 1P).

### Blocking TGFβ signaling, p38 MAPK activation, and the ELAV-ARE interaction during Trial 2 blocks the ApELAV-*apc/ebp* mRNA interaction

Our hypothesis, that ApELAV is recruited by Trial 2 and binds to the *apc/ebp* transcript, predicts that the ApELAV-*apc/ebp* mRNA interaction should be increased at the 1 hr time point following Trial 1 in ganglia which receive two-trial training. To directly test this hypothesis, we performed a cross-linking immunoprecipitation experiment followed by qPCR (CLIP-qPCR), to compare the physical interaction of *apc/ebp* mRNA with ApELAV between experimental (two-trial training) and within-animal control groups (Figure 3a_1_). Following two-trial training, we indeed found a significant increase in *apc/ebp* mRNA association with ApELAV (Figure 3a_2_; Table 1Q), which strengthens the hypothesis that ELAV, by binding to *apc/ebp* transcript, promotes its stabilization at that timepoint.

**Figure 3.**
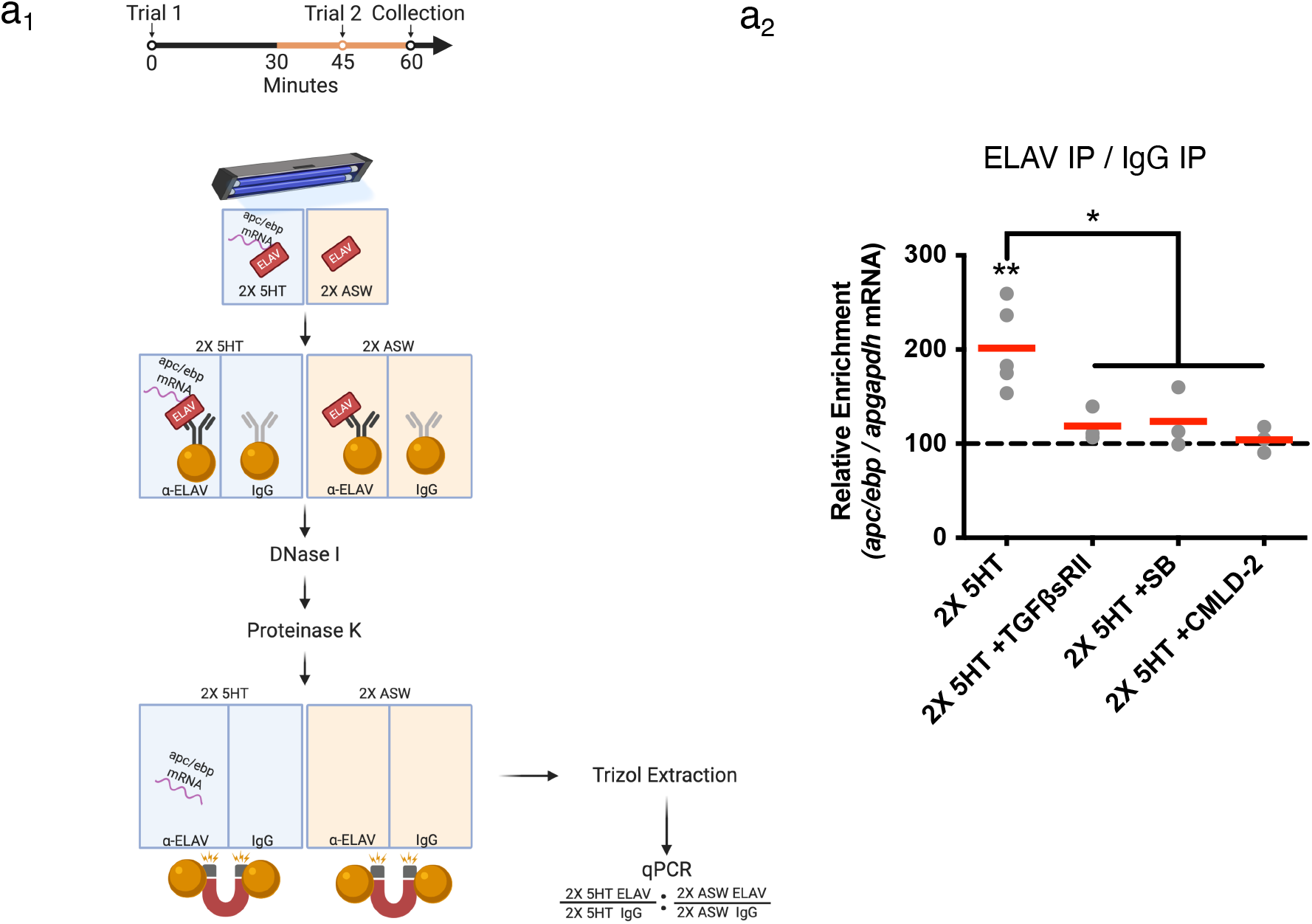
Cross-linking immunoprecipitation (CLIP)-qPCR reveals increase in ApELAV-*apc/ebp* interaction following two-trial training. **(a) (Top)** Experimental paradigm. SN somata are UV-fixed and collected at 1 hr following the onset of Trial 1. Dissected ganglia are treated with CMLD-2, SB203580, or TGFβsRII from 15 min before Trial 2 until collection. **(Bottom)** CLIP-qPCR experimental flowthrough. See Methods for more details. **(b)** Following two-trial training, there is a significant increase in ApELAV-*apc/ebp* mRNA interaction which is blocked by treatment with TGFβ-sRII, SB203580, or CMLD-2. Each data point represents the average of 3 full experiments of SN clusters pooled from 8 animals. \**p*<0.05, \*\**p*<0.01; n=3-5

If ApELAV is responsible for stabilizing *apc/ebp* mRNA during Trial 2, thereby prolonging its expression to 1 hr following Trial 1, then blocking TGFβ signaling should block the ApELAV-*apc/ebp* mRNA interaction. We have previously shown that treatment with TGFβsRII disrupted the increase in Trial 2-dependent *apc/ebp* mRNA level (Kopec et al., 2015). Treatment with TGFβsRII resulted in the complete loss of the ApELAV-*apc/ebp* mRNA interaction at the same time point when assayed by CLIP-qPCR and compared to contralateral within-animal control ganglia which received only vehicle (0.1% BSA in ASW) (Figure 3a_2_; Table 1R), demonstrating that TGFβ signaling is required for the increase in ApELAV-*apc/ebp* mRNA interaction at that time point.

Activation of p38 MAPK is dependent on intact TGFβ signaling during Trial 2 (Figure 2a). Thus, we predicted that blocking p38 MAPK should block the ability of ApELAV to interact with *apc/ebp* mRNA. We found that blocking p38 MAPK with SB203580 blocked the ApELAV-*apc/ebp* mRNA interaction compared to contralateral within-animal control ganglia which received only vehicle (0.1% DMSO) (Figure 3a_2_: Table 1S). These results demonstrate that TGFβ signaling and p38 MAPK activation during Trial 2 are required for the interaction of ApELAV and *apc/ebp* mRNA.

To directly demonstrate if CMLD-2 blocks the ApELAV-*apc/ebp* mRNA interaction, we treated ganglia with CMLD-2 during Trial 2 and performed CLIP-qPCR. We found that treatment with CMLD-2 during Trial 2 significantly disrupted the ApELAV-*apc/ebp* mRNA interaction compared to contralateral within-animal control ganglia which had received only vehicle (Figure 3a_2_; Table 1T). These results demonstrate the effectiveness of CMLD-2 as an inhibitor of the ApELAV-*apc/ebp* mRNA interaction in our two-trial paradigm.

### CMLD-2 disrupts the induction of LTM by Trial 2

The molecular observations described thus far provide clear predictions for the role of TGFβ-mediated post-transcriptional regulation of *apc/ebp* mRNA by the RNA-binding protein ApELAV in the behavioral induction of LTM. We have previously shown that TGFβ signaling during Trial 2 is required for (i) the expression of *apc/ebp* mRNA at the 1 hr time point and (ii) the induction of LTM for sensitization (Kopec et al., 2015). In the present paper, our data show that TGFβ signaling during Trial 2 provides stabilization of *apc/ebp* mRNA by means of increasing its association with ApELAV. Thus collectively, our molecular data predict that the ApELAV-*apc/ebp* mRNA interaction during Trial 2 is required for the prolonged increase of *apc/ebp* mRNA that is necessary for the induction of LTM. We directly tested this prediction in a final set of behavioral experiments examining LTM for sensitization of the tail-elicited siphon withdrawal reflex (T-SWR). We used the T-SWR semi-intact preparation (Sutton et al., 2001; Kopec et al., 2015) that permits the manipulation of the molecular environment of the CNS while directly assaying withdrawal responses (see Figure 4a_1_, Methods).

**Figure 4.**
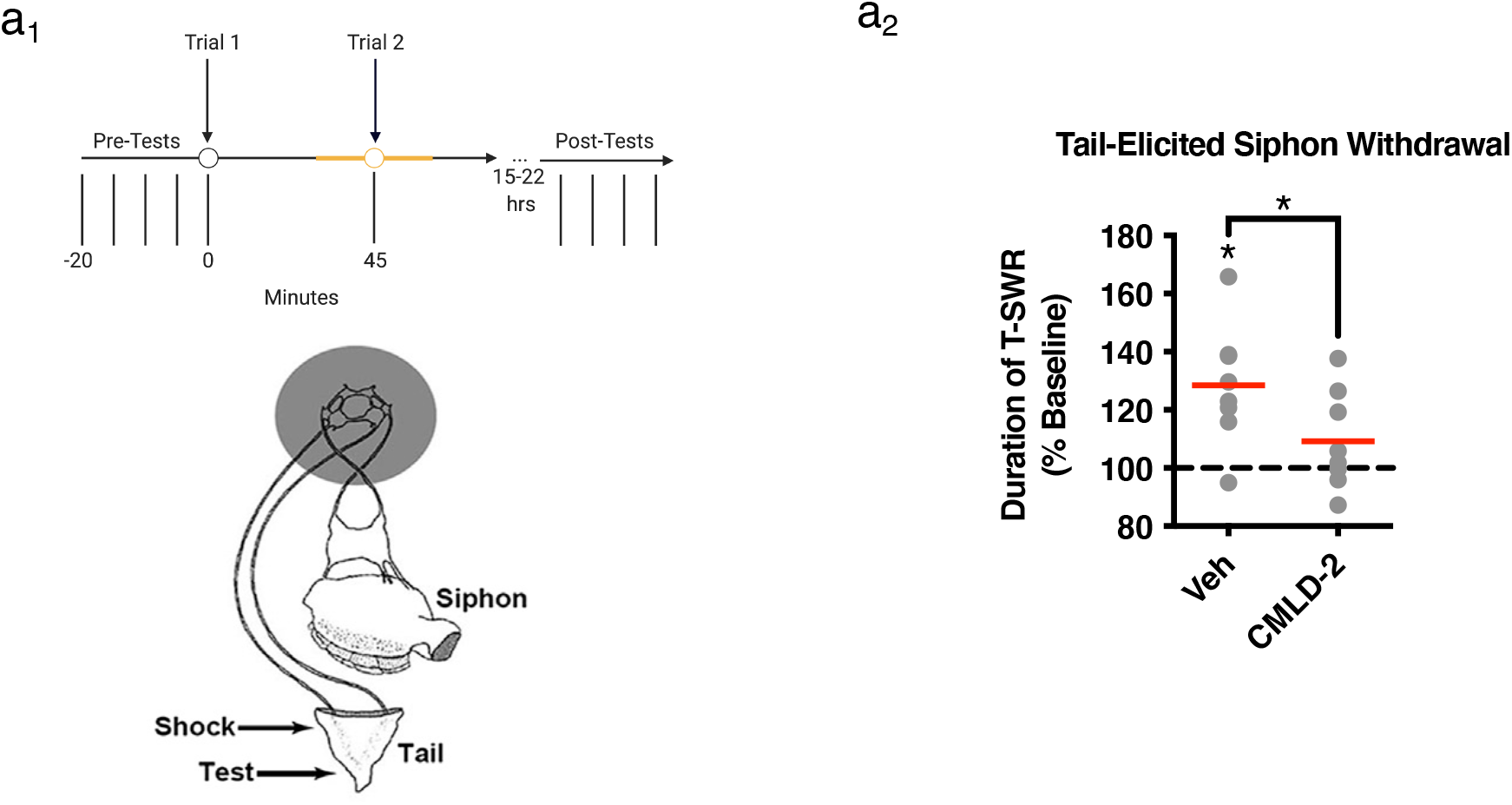
ApELAV-*apc/ebp* mRNA interaction is required for LTM. **(a_1_) (Top)** Experimental paradigm. LTM is assessed by stimulating the test site before training, then 15-22 hr after training and measuring the T-SWR. **(Bottom)**Semi-intact preparation. Two-trial behavioral training is administered to the training site, and drug is applied to the isolated CNS chamber. **(a_2_)** Blocking the ApELAV-*apc/ebp* mRNA interaction during Trial 2 significantly disrupts LTM formation. \**p*<0.05; n=8.

To test the hypothesis that the ApELAV-*apc/ebp* mRNA interaction is required during Trial 2 for LTM formation, we exposed the CNS to 100 μM CMLD-2 or an equivalent volume of vehicle during Trial 2 (Figure 4a_1_). In the presence of vehicle, a within-group comparison revealed significant LTM for sensitization of the T-SWR. In contrast, the induction of LTM was significantly disrupted when the ApELAV-*apc/ebp* mRNA interaction was blocked during Trial 2. Moreover, a between-group comparison revealed a significant difference between vehicle and CMLD-2 groups (Figure 4a_2_; Table 1U). These behavioral data confirm the predictions derived from our molecular observations and support a general model in which TGFβ signaling in Trial 2 mediates RNA stabilization that is essential for LTM formation.

## Discussion

Our results reveal *with temporal precision* a novel molecular mechanism underlying LTM formation: the stabilization of previously synthesized mRNA by a repeated training trial. Specifically, we show that *apc/ebp* mRNA is post-transcriptionally stabilized by the RNA-binding protein ELAV by a repeated training trial. This stabilization is dependent upon TGFβ signaling and p38 MAPK activity, which is required for the induction of LTM (Kopec et al., 2015). Our data supports a model (Figure 5) in which Trial 1 gives rise to TrkB signaling, resulting in an increase in CREB-mediated transcription and an increase in *apc/ebp* gene expression 45 min later. During Trial 2, TGFβ-like ligands act in SNs to activate ELAV through p38 MAPK causing an increase in its binding to *apc/ebp* mRNA, thereby stabilizing the transcript, resulting in a transcription-independent prolonged increase in *apc/ebp* mRNA level. Finally, the ApELAV-mRNA interaction during Trial 2 is a pre-requisite for LTM formation. These experiments provide novel evidence for a GF-mediated process in LTM formation resulting from post-transcriptional regulation that modulates the stability of an immediate early gene transcript, which serves as a transcription factor required in LTM.

**Figure 5.**
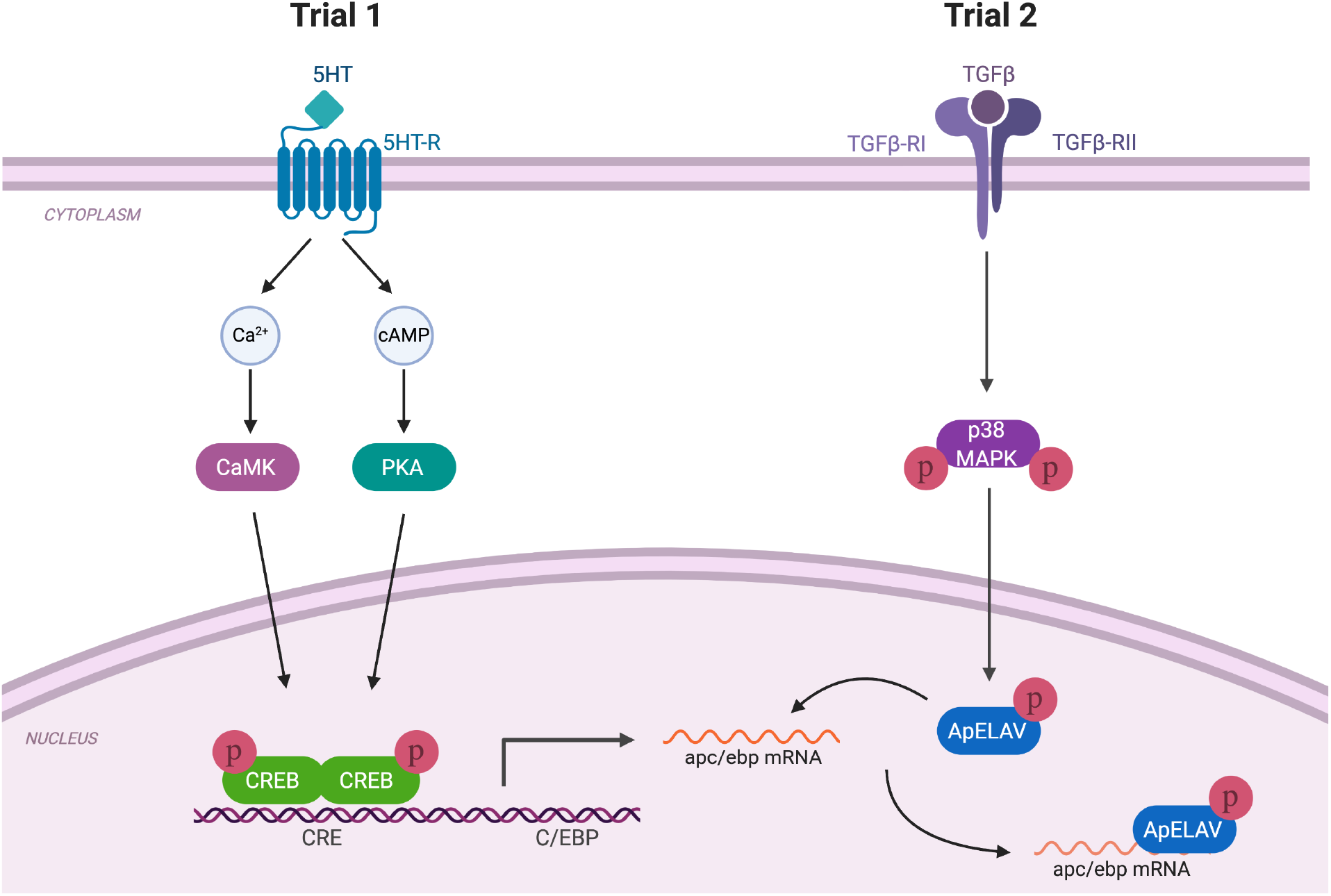
Working model of molecular mechanisms underlying *apc/ebp* transcript stabilization in two-trial LTM formation. Trial 1 gives rise to an increase in CREB mediated transcription and an increase in *apc/ebp* gene expression 45 min later. When Trial 2 occurs at that timepoint (45 min), TGFβ signaling gives rise to an increase in p38 MAPK activation, resulting in an increase in ELAV-*apc/ebp* interaction. This interaction increases the stability of *apc/ebp* transcript and prolongs its expression to 1 hr, which is permissive for the induction of LTM.

### C/EBP in Long-Term Memory Formation

The C/EBP family of transcription factors (having 6 isoforms) are immediate early genes downstream of CREB (Alberini, 2009). C/EBPs, particularly the C/EBPβ and C/EBPδ isoforms, are regulated by learning-related stimuli across a wide range of species, as well as across diverse brain regions and learning tasks (Alberini et al., 1994; Alberini, 2009). Together with these studies, our findings indicate that C/EBP signaling is a significant molecular step mediating plasticity and memory formation. Interestingly, C/EBP contributes to the induction of GF ligands and receptors in (i) IGF-II in hippocampus (Chen et al., 2011), (ii) VEGF-C and its receptor in lymphatic endothelial cells (Min et al., 2011), (iii) TGFβr-II in human embryonic stem cells (Takayama et al., 2014), and (iv) NGF in cerebral cortex (McCauslin et al., 2006). This raises the intriguing possibility of a GF-mediated positive feedback loop during memory formation in which GF signaling upregulates C/EBP, which in turn increases GF ligand and/or receptor expression.

Here we have observed two phases of *apc/ebp* gene expression during LTM formation with two-trial training: First, following Trial 1, a *transcription-dependent* increase in *apc/ebp* mRNA level, and second, following Trial 2, a *transcription-independent* stabilization via the RNA-binding protein ApELAV, which prolongs the increase in *apc/ebp* mRNA level. This is a critical point of regulation since the presence of *cis*-acting regulatory elements on specific mRNAs, such as AREs, can modulate, or even counteract, the effect of increased transcription (Ross, 1995; Lee et al., 2015). The AU-rich element RNA binding proteins ApAUF1 and ApELAV can bi-directionally regulate *apc/ebp* mRNA stability: ApAUF1 binding to the 3’UTR of *apc/ebp* induces the degradation of the transcript, and overexpression of ApAUF1 inhibits 5HT-induced LTF in SN-MN co-culture (Lee et al., 2012). Conversely, ApELAV binds to the same AU rich domain but stabilizes the transcript (Yim et al., 2006). Using the small-molecule inhibitor CMLD-2 to block the ApELAV-*apc/ebp* mRNA interaction during the second trial of two-trial training, we demonstrated both the block of the increase in *apc/ebp* mRNA level and the disruption in the induction of LTM, underscoring the importance of this stabilizing mechanism for prolonging mRNA level and its critical role in LTM formation. Thus, the expression of ApAUF1 and ApELAV, or the ratio of these proteins, has the capacity to favor either the degradation or stabilization of *apc/ebp* mRNAs.

Prior studies have demonstrated a potential role for ELAV RNA-binding proteins in LTM formation. The present data significantly extend these initial studies in two ways: They demonstrate exactly *when* during LTM formation ELAV proteins are engaged, and they elucidate a *mechanistic link* between GF signaling, mRNA stabilization, and LTM formation. Specifically, in previous studies, the expression of the mammalian neuron-specific ELAV protein HuC has been shown to increase in mouse hippocampus following LTM training, and this increase was concomitant with an increase in its target mRNA GAP-43. Additionally, knockdown of HuC in mouse hippocampus resulted in impaired spatial learning (Quattrone et al., 2001). Other studies demonstrated that the expression of the mammalian neuron-specific ELAV protein HuD is increased following contextual fear conditioning in mice, and transgenic overexpression resulted in deficits in associative and spatial learning (Bolognani et al., 2004; Bolognani et al., 2007). Collectively, these results demonstrate an important role for ELAV proteins in LTM formation. However, they also pose potentially conflicting views of ELAV’s actions, since both knocking out ELAV (Quattrone et al., 2001) and overexpression of ELAV (Bolognani et al., 2007) leads to memory deficits. The conflicting nature of these findings may be due to an experimental lack of temporal specificity. Using the two-trial paradigm in *Aplysia*, we are able to investigate mRNA regulation on the timescale of minutes. We find that ELAV target mRNA stabilization occurs without any appreciable change in ApELAV protein level, and the interaction between ApELAV and *apc/ebp* mRNA is significantly increased following Trial 2. These findings allowed us to investigate the specific mechanistic link between a single, unique training stimulus (Trial 2), and mRNA stabilization, which we further demonstrate occurs through p38 MAPK activation downstream of TGFβ signaling.

Finally, we should emphasize that, in addition to transcript stabilization, other mechanisms of post-transcriptional regulation have previously been implicated in LTM formation in *Aplysia* (Rajasethupathy et al., 2009).

### TGFβ signaling and RNA-binding proteins

Signaling through distinct GF families is specifically engaged, both spatially and temporally, during LTM formation (Kopec et al., 2015). In the first trial of a two-trial training paradigm, TrkB signaling at the SN synapse is required for *apc/ebp* gene expression 45 min following the onset of the trial, and TGFβ signaling is required at the SN soma during the second trial (at 45 min) for prolonged increase in *apc/ebp* mRNA level. Here, we demonstrate that: (i) direct activation of TGFβ signaling (in the absence of 5HT at 45 min) results in a transcription-independent increase in *apc/ebp* mRNA level, and (ii) this effect is generated by recruiting ApELAV to bind and stabilize the *apc/ebp* transcript produced by TrkB signaling-dependent transcriptional mechanisms initiated by Trial 1.

Interestingly, there is considerable evidence that TGFβ-1 initiates stabilization of mRNAs via an RNA binding complex (Amara et al., 1995). In cardiac fibroblasts, TGFβ-1 treatment induced HuR (a member of the mammalian ELAV family) translocation from the nucleus to the cytoplasm where it bound the ARE region of the 3’ UTR in TGFβ-1 mRNA thus stabilizing the mRNA and increasing protein level (Bai et al., 2012). These data suggest the intriguing possibility that, in addition to TGFβr-II signaling initiating a cascade to prolong the expression of critical learning-related genes like *apc/ebp*, other GF ligands could be induced and stabilized in the same way (including a TGFβ-1-like ligand itself). Indeed, GF ligands (Lim and Alkon, 2012) as well as their receptors (Balmer et al., 2001) can be regulated by ARE-RNA binding proteins.

The mammalian ELAV-like protein HuR is phosphorylated by PKC and p38 MAPK (Doller et al., 2007; Doller et al., 2008; Lafarga et al., 2009; Doller et al., 2011; Eberhardt et al., 2012; Mirisis and Carew, 2019). In our two-trial paradigm, we found that p38 MAPK activity was engaged through TGFβ, and its activity was required both for the ApELAV-*apc/ebp* mRNA interaction and for the prolonged increase in *apc/ebp* mRNA. Indeed, phosphorylation of ELAV in many systems has been shown to (i) increase its binding affinity for AREs (Pascale et al., 2005; Doller et al., 2011; Eberhardt et al., 2012), (ii) modulate the binding affinity of ELAV to specific types of AREs (Doller et al., 2010; Eberhardt et al., 2012), and (iii) regulate its nucleo-cytoplasmic shuttling ability (Fan and Steitz, 1998; Wang et al., 2002; Doller et al., 2008; Bai et al., 2012; Doller et al., 2015). Interestingly, p38 MAPK activation has additionally been shown to inhibit the activity of destabilizing ARE-binding proteins, which would act to shift the balance towards mRNA stabilization (Briata et al., 2005).

In conclusion, the present results, taken with previous work, illustrate the importance of transcript regulation through GF signaling as a critical site of molecular interaction underlying long-term memory formation.

## AUTHOR CONTRIBUTIONS

AAM conceived of, designed, performed, and analyzed experiments. AMK conceived of, designed, performed, and analyzed experiments. TJC conceived of and designed experiments. All authors wrote the paper.

## ACKNOWLEDGEMENTS

We thank A. Alexandrescu, P.E. Miranda, and N.V. Kukushkin for their comments on the manuscript. We thank Bong-Kiun Kaang for sharing ApELAV plasmids. We thank the Hochwagen Laboratory for use of the Deltavision Elite microscope for smFISH imaging and the NYU Center for Genomics and Systems Biology Genomics Core for use of the Roche 480 LightCycler for qPCR experiments. Figures were created with BioRender.com. This work was supported by NIMH R01 MH 094792 to TJC and a Hellenic Medical Society of New York Leonidas Lantzounis Research Grant to AAM.

---

**Supplementary Figure 1.**
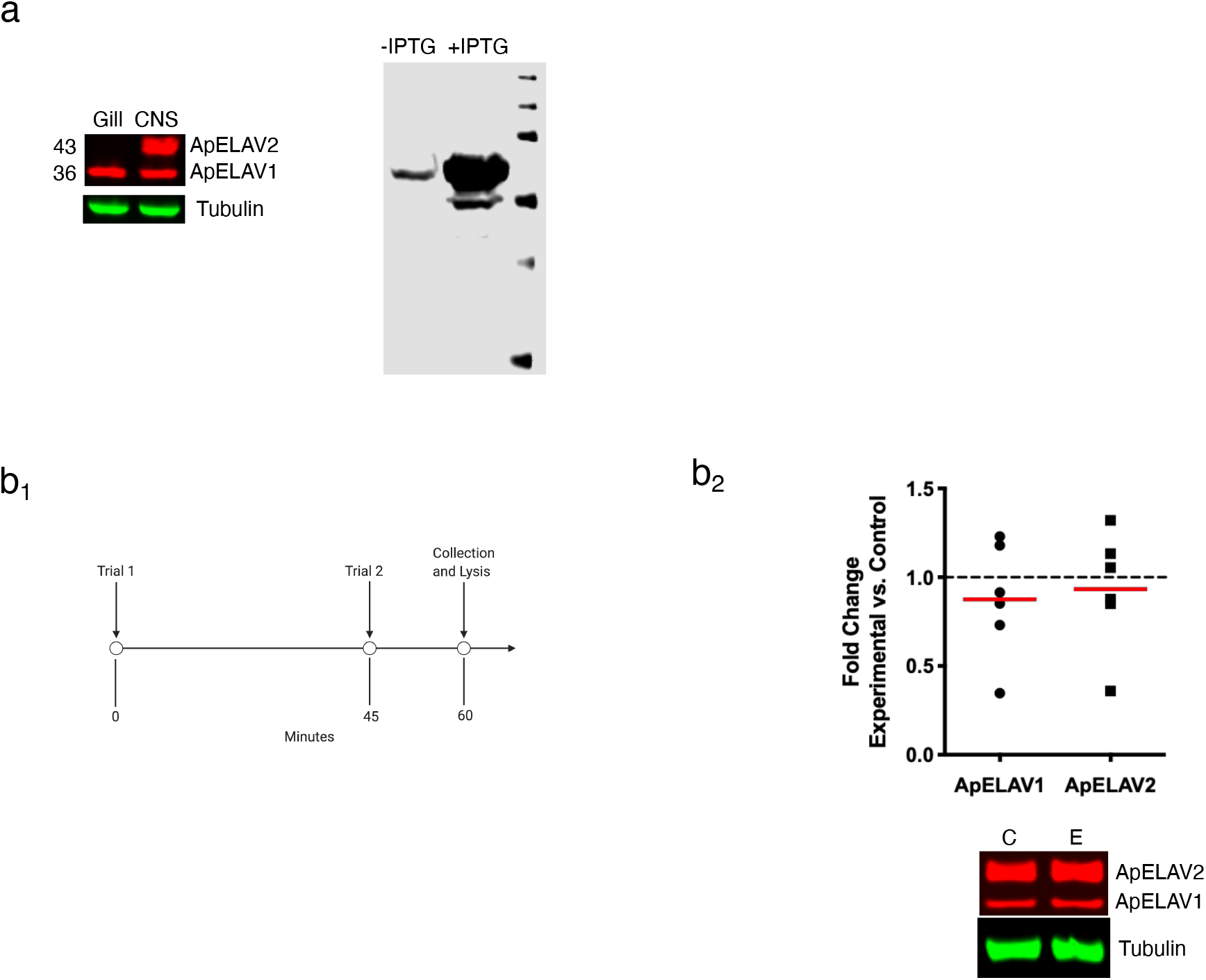
**(a) (Left)** ApELAV1 and ApELAV2 are recognized by the antibody at their respective predicted molecular weights. ApELAV1 is ubiquitously expressed and ApELAV2 is neuron-specific. **(Right)** The antibody reacts strongly with recombinant ApELAV1 (expression induced with IPTG). **(b_1_)** Experimental paradigm. Experimental ganglia receive two-trial training with 5HT and contralateral control ganglia receive two-trial training without 5HT. Ganglia are collected and lysed at 1 hr. **(b_2_)** There is no significant change in ApELAV protein levels following two-trial training. Representative western blots are shown below the histograms (*n*=6).

**Supplementary Figure 2.**
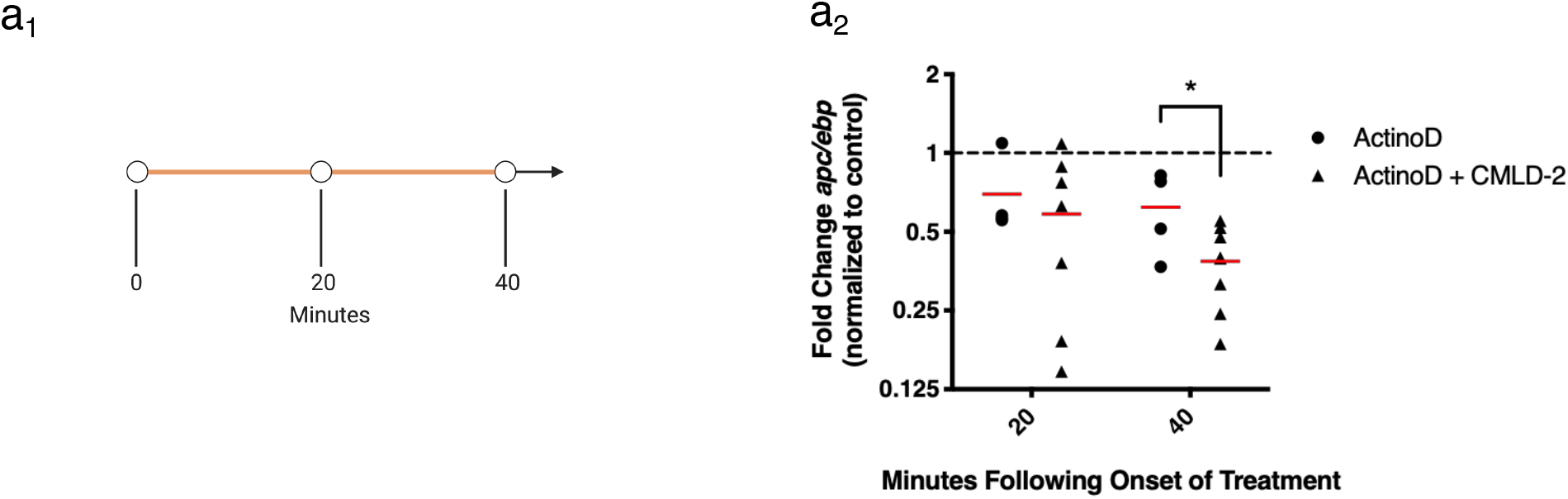
**(a_1_)** Experimental paradigm. Experimental ganglia are treated with actinomycin D and CMLD-2, or actinomycin D alone, and control ganglia are treated with vehicle. SN clusters are collected and lysed at 20 min or 40 min following treatment onset. (a_2_) CMLD-2 treatment results in significantly lower *apc/ebp* levels at 40 min post-treatment onset in experimental ganglia compared to actinomycin D alone.

**Supplementary Figure 3.**
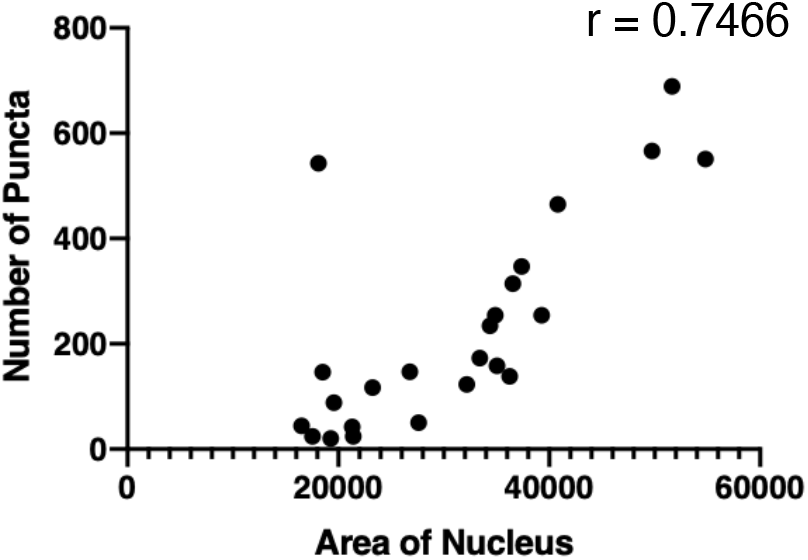
Distribution of smFISH data from 2X ASW +DMSO (control) group. There is a strong correlation between cell size (measured by area of nucleus) and number of puncta observed.

**Supplementary Table 1.**
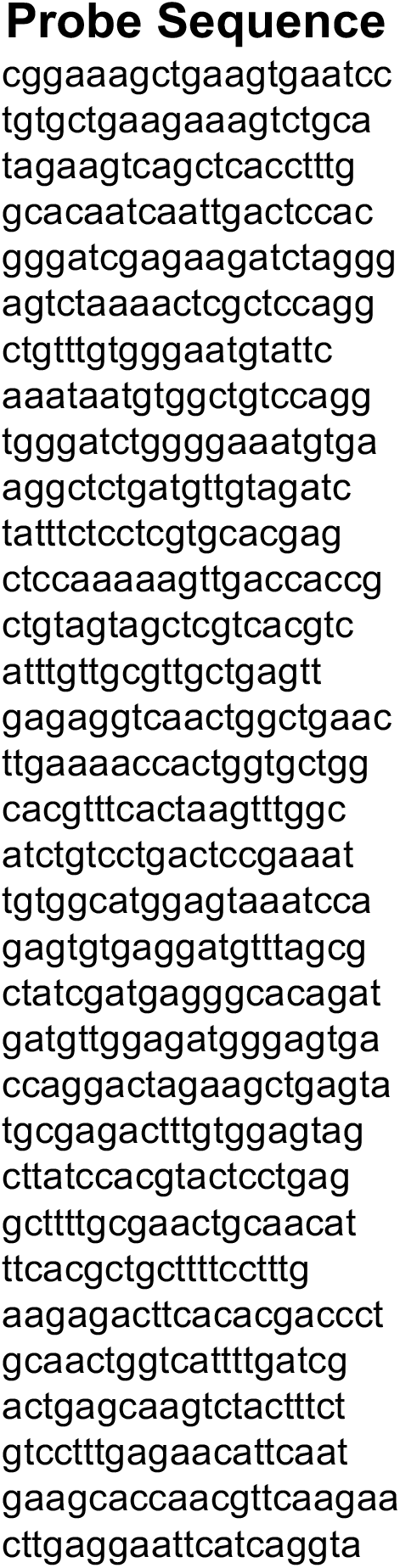
List of DNA probes custom designed by LGC Biosearch Technology for detection of *apc/ebp* mRNA.

